# An In Silico Model to Simulate the Evolution of Biological Aging

**DOI:** 10.1101/037952

**Authors:** A. Šajina, D.R. Valenzano

## Abstract

Biological aging is characterized by an age-dependent increase in the probability of death and by a decrease in the reproductive capacity. Individual age-dependent rates of survival and reproduction have a strong impact on population dynamics, and the genetic elements determining survival and reproduction are under different selective forces throughout an organism lifespan. Here we develop a highly versatile numerical model of genome evolution — both asexual and sexual — for a population of virtual individuals with overlapping generations, where the genetic elements affecting survival and reproduction rate at different life stages are free to evolve due to mutation and selection. Our model recapitulates several emerging properties of natural populations, developing longer reproductive lifespan under stable conditions and shorter survival and reproduction in unstable environments. Faster aging results as the consequence of the reduced strength of purifying selection in more unstable populations, which have large portions of the genome that accumulate detrimental mutations. Unlike sexually reproducing populations under constant resources, asexually reproducing populations fail to develop an age-dependent increase in death rates and decrease in reproduction rates, therefore escaping senescence. Our model provides a powerful *in silico* framework to simulate how populations and genomes change in the context of biological aging and opens a novel analytical opportunity to characterize how real populations evolve their specific aging dynamics.

## I. INTRODUCTION

Natural populations evolve from the interaction between external selective forces and the internal capacity to respond to them. External forces include predators, parasites and available resources, and population response to these forces importantly depends on rates of mutation and recombination. Standard evolutionary theory of aging predicts that deleterious mutations affecting early or late survival are differentially selected, with mutations negatively affecting survival in early life being readily removed by purifying selection, as opposed to those affecting survival in late life, which instead have a tendency to accumulate due to weaker selective forces^1,5,6,9,11^. A consequence of this phenomenon is that young individuals have lower death rates (i.e. higher chances to survive) compared to old individuals. Furthermore, it is theoretically possible that selection could favor alleles that have a detrimental effect on survival in late life stages, as long as they have a beneficial effect early on in life – i.e. alleles that contribute to an aging phenotype can be fixed in the gene pool if their overall effect on fitness is positive^12^.

Although these models greatly helped to frame the genetics of biological aging on a solid evolutionary basis, there are some important aspects that were left out, which are key for our understanding on how selection shapes individual fitness in biological populations. In particular, exclusively focusing on how selection shapes age-specific survival, they did not analyze how genes that regulate survival and reproduction might coevolve and how their interaction could affect individual fitness and population stability under different selective pressures, which are key aspects to understanding how natural populations evolve.

Unlike analytical methods, computer simulations – or numerical models – can sustain high parameter complexity and are particularly suited to model the evolution of dynamic systems, such as biological systems^2,10^.

To analyze the role of selection in shaping the genetic basis of reproductive aging and age-dependent changes in death rate, we developed an *in silico* model that simulates the evolution of biological populations, where each individual agent is provided with a genome that defines the probabilities to reproduce and survive. We let a genetically heterogeneous population evolve under different environmental constraints, including limiting resources, sexual or asexual reproduction and different population sizes. We find that, in a sexual model, the force of selection decreases with age after sexual maturation. However, we demonstrate that, in stable environments, selection favors prolonged survival and reproduction after sexual maturation more than in unstable conditions. In unstable environments, characterized by continuous expansions and compression of population size, we observe the emergence of increased early mortality. Importantly, we observe that the hallmarks of biological aging, i.e. the time-dependent decrease in individual survival and reproduction, are more dramatic in the sexual model and do not evolve in the asexual model under constant resource conditions. Our model offers the possibility to address key evolutionary questions regarding the evolution of biological aging at both individual and population level, and its applications can give key insights on how genomes contribute to the time-dependent deterioration of biological functions.

## II. THE MODEL

### a. The genome

Our model simulates the evolution of a population of agents, which evolves through a sequence of discrete time intervals, called *stages.* For each agent *i*, the probability to surviving and reproducing at each stage is defined by a bit-string code that we name genome (*G_i_*). The probability to surviving from one stage to the next is indicated by *s_k_*, and we refer to it as *transition probability.* The transition probability for stage k is proportional to the numbers of 1’s present in a corrensponding 20-bits array, called *S_k_* (FIG. 1). Analogously, the probability to reproducing at stage k, indicated by *r_k_*, is proportional to the number of 1’s in the 20-bits array *R_k_*:

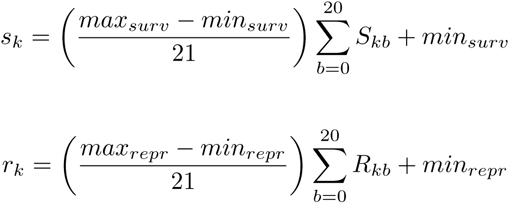

with both *S_kb_* and *R_kb_* that can be = 0 or 1 and *max_surv_* and *max_repr_* being the initially set maximum allowed survival and reproduction probability, respectively.

**FIG. 1.**
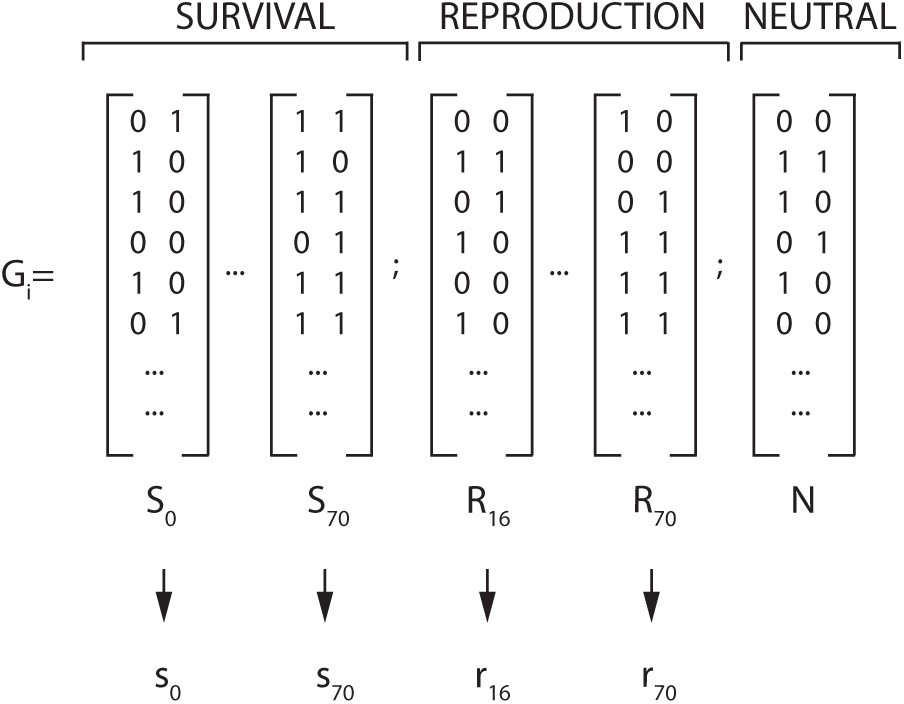
Genome composition. The genome *(G_i_)* contains a sequence of discrete bit arrays – *S_i_* and *R_i_* – that define the probability to survive from one stage to the next and to reproduce at each stage, respectively indicated as *s_i_* and *r_i_*. Reproduction starts at age 16, therefore there are no *R* and *r* values with an index smaller than 16. A neutral portion of the genome, indicated by *N*, has no corresponding phenotype, and is used as a control region to measure neutral evolution.

*G_i_* also contains a bit array that is not translated in any phenotype (indicated as *N* in FIG. 1), which is used as a functionally-neutral portion of the genome.

For each agent *i* in the population, *G_i_* is a continuous bit array, i.e. all *S_k_*, *R_k_* and *N* are contiguous. For representation purposes, in all the plots shown in results, both *s_k_* and *r_k_* are represented in ascending order, i.e. with *s_k_* preceeding *s_k+1_*. However, the actual order along the bit array is randomized, so that *S_k_* does not necessarily preceed *S_k+1_*.

### b. Simulation progression

The seed (or starting) population is composed of genetically heterogeneous agents, whose genomes are randomly generated bit arrays, which have on average an equal number of 1’s and 0’s. Each agent from the seed population has a specified chronological age, and becomes one time-unit older at every new stage. Resources are used as the population growth-limiting factor. Resource amount is defined before the simulation starts and, at each stage, the population can be in a state of resource abundance or shortage, i.e. *N_t_* ≤ *R_t_* and *N_t_* > *R_t_*, with *R_t_* being resources available at stage *t* and *N_t_* being the populations size (i.e. number of individuals) at stage *t*. In the former situation, for each individual, the death rate will correspond to 1 − *s_t_*, while in the latter it will correspond to *µ* (1 − *s_t_*), with *µ* being a constant that increases actual death rate for all the agents in case of resource shortage. Depending on the simulation type, resources are set to be either constant or variable. In the *constant resource* model, resources are the same at every new stage, regardless from the leftover resources from the previous stage. However, population growth and contraction will affect the death rate depending on whether population size exceeds resouce units. In the *variable resource* model, the amount of available resources is the sum of two quantities. One is proportional to the difference between resources and population size from the previous stage (leftover resources), and the other is a fixed resource increment 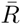:

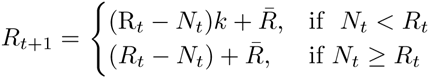

If the amount of resource units from the previous stage exceeds population size, the leftover resources regenerate proportionally to a fixed proliferating value *k* – set at the beginning of the simulation – and are added to the fixed resource increment 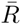. However, if the population size exceeds or is equal to the resource units, i.e. no resource is left from the previous stage, the difference between population size and resources available is subtracted from the fixed resource increment 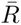. If this difference is ≤ 0, resources are then set to 0. In this case, death rate is affected (see above).

Reproduction follows resource consumption (FIG. 2). In the asexual model, each individual can reproduce or not, based on the probability provided by the individual age-specific *r_k_* value. In the sexual model, the individuals that, based on the stage-specific *r_k_* value, are selected to reproduce, undergo chromosome recombination. Recombination is determined by a fixed recombination frequency, and is followed by the random selection of one of the two chromosomes (either the left or the right in each *S*_0:70_, *R*_16:70_, and *N* from FIG. 1) to match a corresponding recombining chromosome selected from another individual. Chromosome matching is followed by mutation, which is set by a fixed mutation rate before each simulation. Mutation transforms 0s to 1s and, vice versa, 1s to 0s. Since any 0 mutating into 1 in both *R_k_* and *S_k_* translates into an increase in the probability of reproducing or surviving, respectively, corresponding to a phenotypically *beneficial* mutation, we let 0 to 1 mutations to be 10 times less likely than 1 to 0 mutations, in order to let mutations to be more likely detrimental than beneficial.

**FIG. 2.**
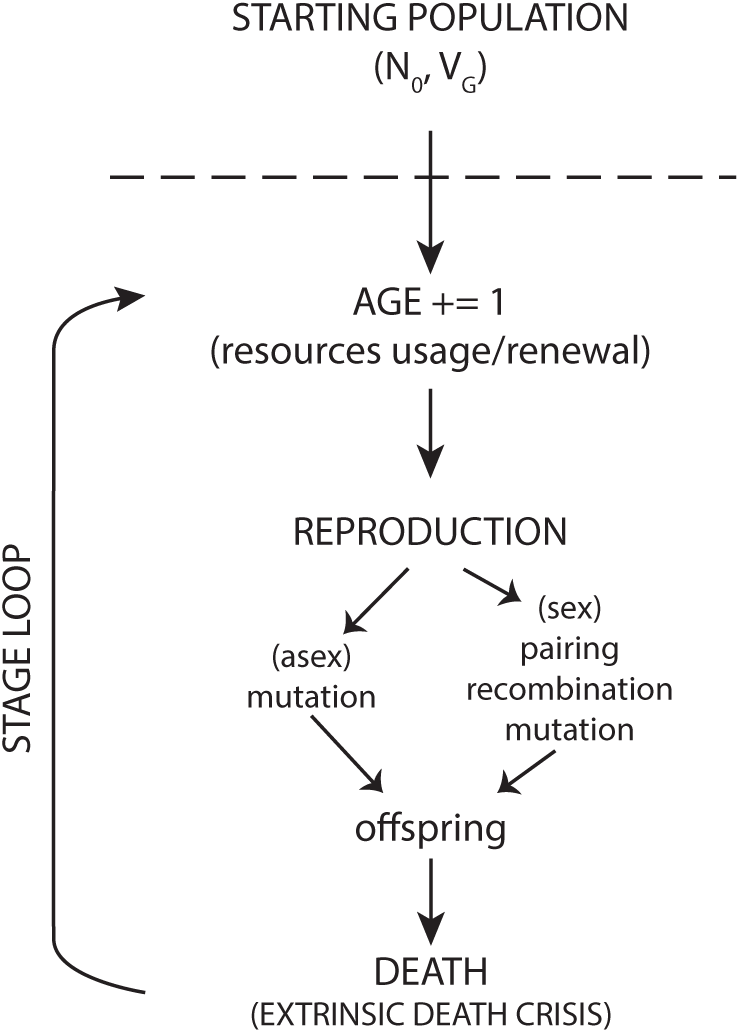
Schematics of events sequence in the simulation. *N*_0_ and *V_G_* represent the initial number of individuals and the genetic variance initially defined for any seed (or starting) population.

Once the *mutation step* takes place, a novel genome is complete, and it represents the new survival and reproduction program for a newborn agent, which is added to the current population with the starting age of 0.

Following reproduction, each agent’s *s_k_* values contribute to determining whether the agent survives or is eliminated from the population. The actual probability of survival through any stage depends on i) each agent’s *s_k_* value for the actual age *k* at the given stage, ii) the difference between resources and population, which can add a weight on each agent’s death risk, and iii) the *extrinsic death crisis* parameter, which can be initially set and determines whether – at specific stages – the population undergoes massive death.

All individuals reach sexual maturation at stage 16, therefore *R_k_* start at *k* = 16.

### A. Fitness

We define individual *genomic fitness* (*GF_ind_*) as the cumulative individual lifetime expected offspring contribution, given the genetically determined probabilities to survive and reproduce. For instance, the *genomic fitness* for an agent at age 50 corresponds to the product of two terms, one that defines the probability to surviving to reproductive age:

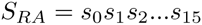

and one that defines the probability to surviving and reproducing at all the stages that follow sexual maturation, up to the last stage (age 50 in this example):

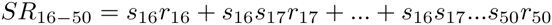

with *s* and *r* as the transition probability for survival and reproduction, respectively, defined in the individual genome by their corresponding *S* and *R* values. More generally, this product can be written as:

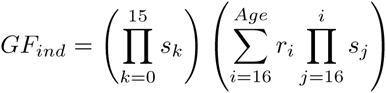

with *Age* as the individual age at which fitness is measured.

We also define, for any stage, an average population genome and its associated *genomic fitness* (*GF_avg_*), as:

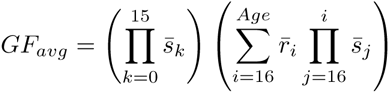

with 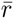 and 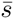 as the average reproduction and survival probabilities for each age.

Analogously, we can define *relative individual genomic fitness* as the ratio of individual fitness to the sum of all the individual fitness:

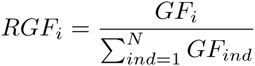

Tracking population size throughout the simulation progression, enables to measure rates of *actual survival* at any stage, for each age class, defined as:

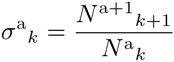

with *N^a^_k_* as the number of individuals of age *k* at stage *a.* Replacing 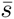 with σ, allows a corrected *GF_avg_*, which we name *population fitness* or *F_p_*:

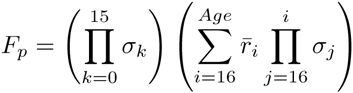

## III. Results

During each simulation, an initial random gene pool is generated, and *S_ik_*, *R_ik_* and *N_i_* (see FIG. 1) evolve freely due to mutation and – in the sexual model – recombination. However, this model also allows populations to be *re-seeded* from end-stages of previous simulations. Population size oscillates as a function of i) available resources (FIG. 3) and ii) evolving distributions of age-dependent reproduction and survival probabilities encoded in each individual genome, which affect the ratios of births over deaths. The model does not bias the direction in which populations will evolve higher or lower survival and reproduction probabilities throughout age, and all the observed changes in age-specific reproduction and survival probabilities (*r_k_* and *s_k_*) are due to selection and drift.

**FIG. 3.**
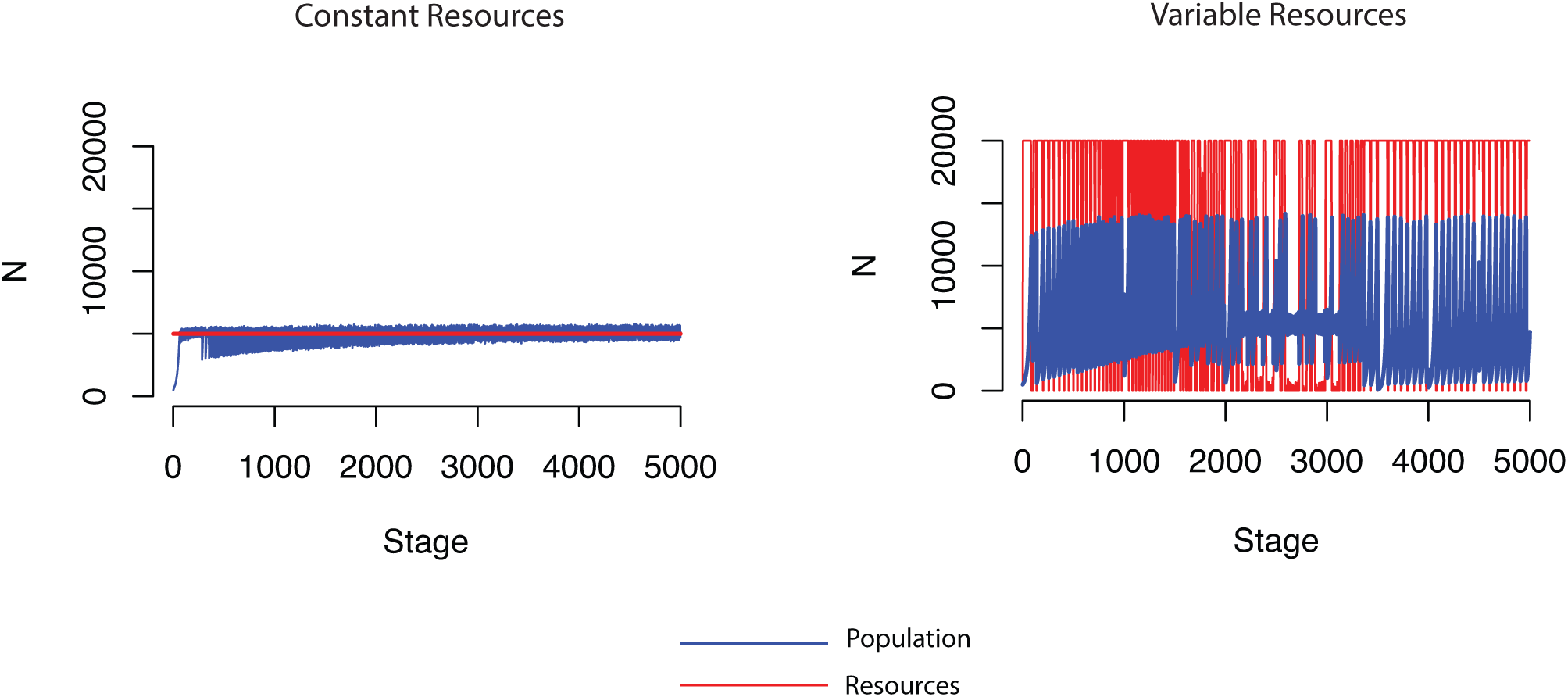
Sample population growth and population size oscillations under constant (left) and variable (right) resource conditions. Only the first 5k stages of the simulations are displayed for simplicity. In the *constant resource* conditions resources are fixed and do not change as a function of population size, whereas, in the *variable resource* conditions, resources vary depending on an intrinsic growth rate, a fixed stagewise increment, and population size.

### A. Sexual model

Under constant resources, populations of agents that reproduce sexually evolve to a state of higher *S_i_* before sexual maturation and continuously declining *S_i_* and *R_i_* values when *i* > *t_m_*, with *t_m_* as the time of sexual maturation (FIG. 4).

**FIG. 4.**
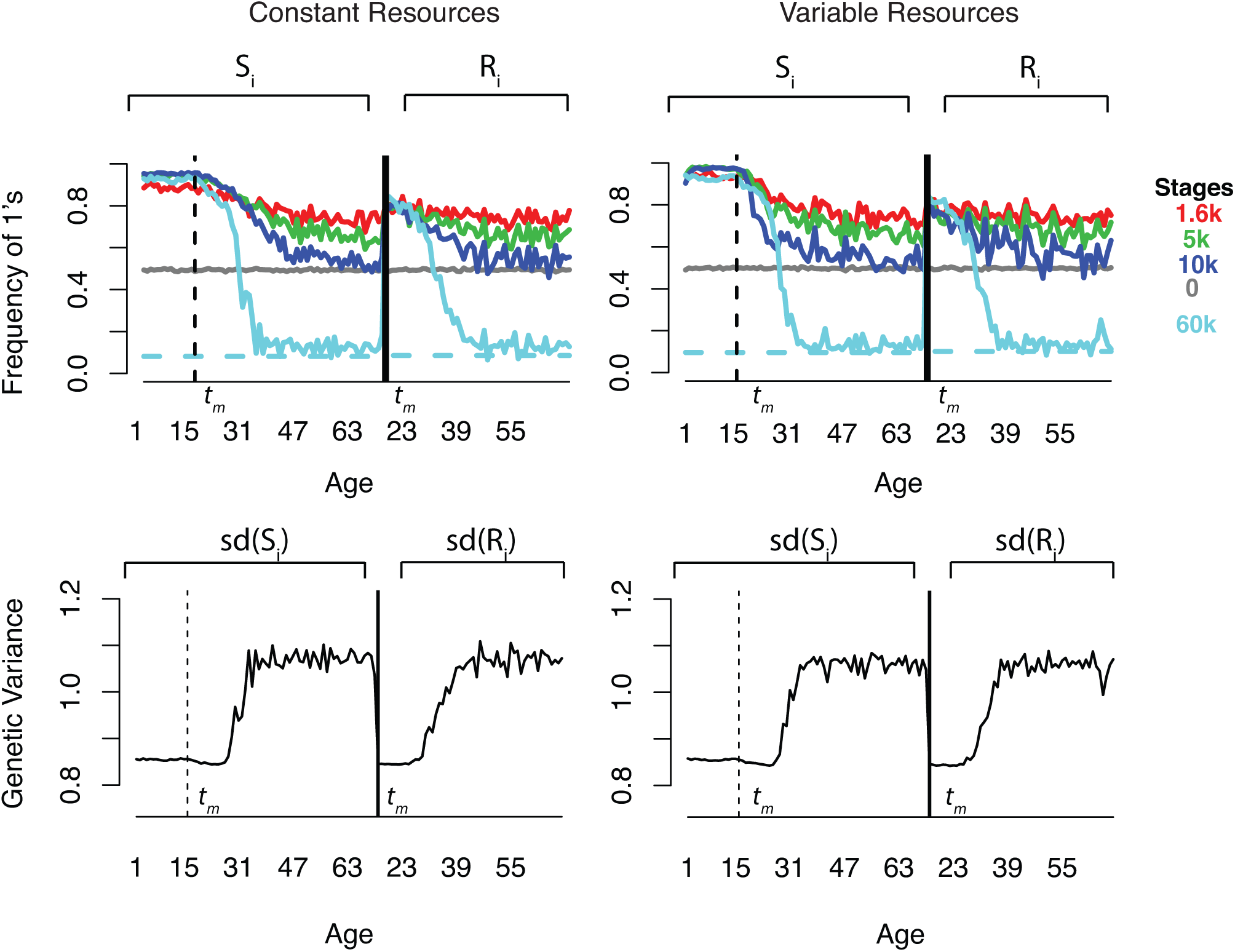
Evolution of the genome in the sexual model. Above: The x-axis represents the position of *S_i_* (on the left) and *R_i_* (on the right) along the genome, sorted according to the age they correspond to. The y-axis represents the ratio of 1’s for each transition probability *S_i_* and *R_i_* indicated on the x-axis. The genomes are randomly generated at stage 0 (grey line) and start with an even distribution of 0’s and 1’s, for all the *S_i_* and *R_i_* (average values only are shown). As the simulation proceeds (different color lines), mutation and selection change the frequencies of 1’s and 0’s for each age-specific *S_i_* and *R_i_*. The genetic modules relative to survival (*S_i_*) fix higher frequencies of 1’s before and about sexual maturation (vertical dashed line and *t_m_*), while reproduction (*R_i_*, not defined before *t_m_*) fix higher frequencies of 1’s starting at *t_m_*. At later stages of the simulation, genetic modules for both survival and reproduction after sexual maturation accumulate more deleterious mutations than at earlier stages, as indicated by the lower values to the right of the cyan curve (60k stages). Below: population average of the genetic variance for *S_i_* and *R_i_* as a function of age after 60k stages of simulation. Genetic Variance before *t_m_* is low, and increases for both survival and reproduction with Age > *t_m_*. Fixed resources are set to 5k units, mutation rate is set to 0.001 per site and recombination rate is set to 0.01.

Since *s_i_* and *r_i_* are directly proportional to *S_i_* and *R_i_,* respectively, also the age-dependent probabilities of surviving and reproducing become accordingly higher in early life and decline after sexual maturation. The phenotypically neutral portion of the genome (indicated by *N* in FIG. 1, and by the dashed lines in FIG. 4 and in FIG. 5B), used as a control region to test the effects of mutation without selection on the genome sequence evolution throughout the simulation, enables to show that the distribution of the *S_i_* and *R_i_* values throughout age is under strong selection before and after sexual maturation, for all the ages where *S_i_* and *R_i_* values are above the curve corresponding to the expected values (horizontal dashed line in FIG. 4 and in FIG. 5B). For each age, the difference between *S_i_*, *R_i_* and the expected values corresponding to the neutrally evolving portion of the genome *(N_i_),* is proportional to the strength of selection.

**FIG. 5.**
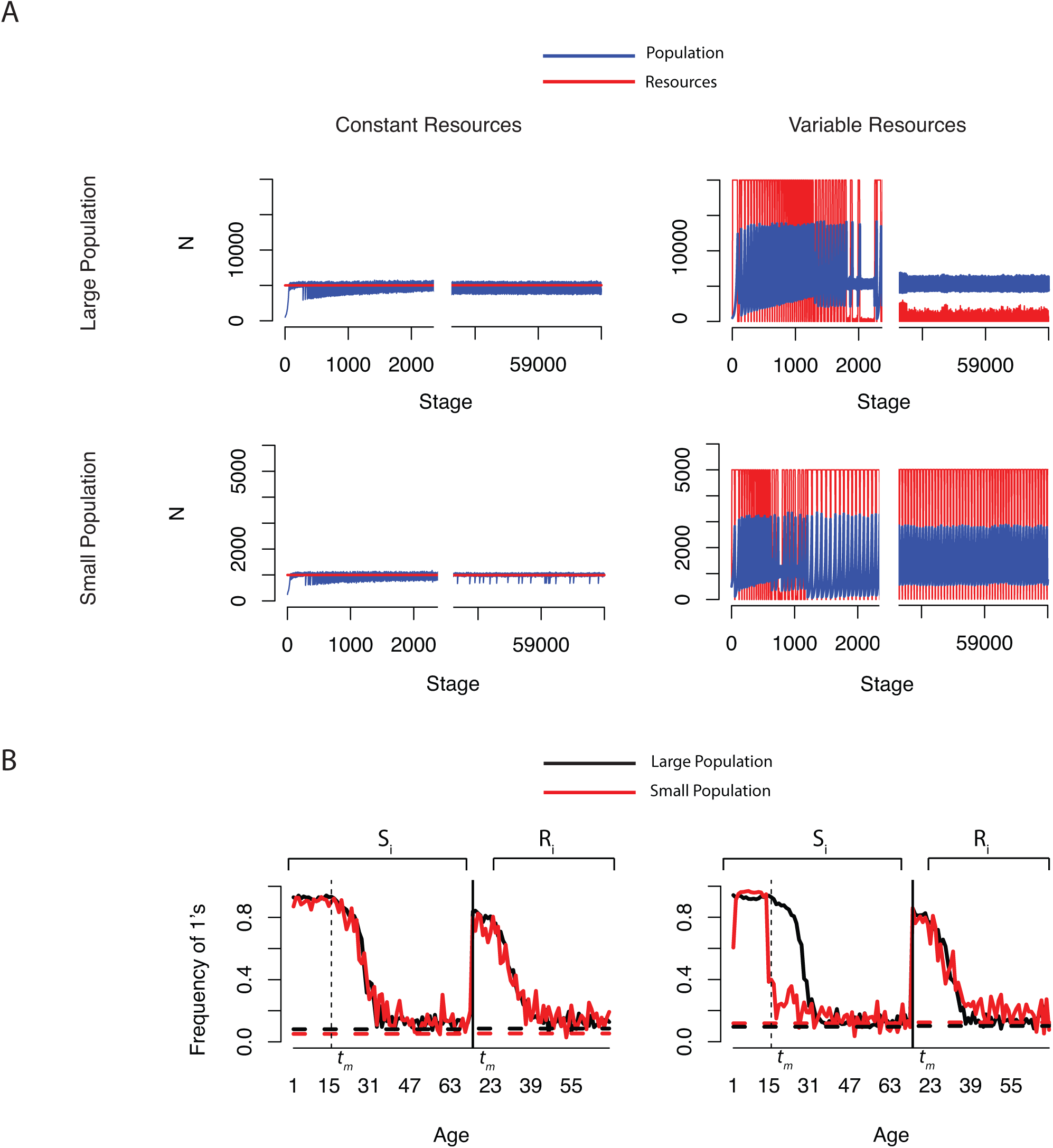
Population, resources and genomic dynamics at different population sizes in the sexual model. A: Population and resource dynamics in larger and smaller populations, under constant (left) or variable (right) resource conditions. Population and resource values are displayed on the y axis. Resources are shown in red and population in blue. Each plot is split in two parts, with early stages on the left, and late stages on the right. Each run consists of 60,000 stages. Large and small populations are determined by the resource value, set to 5k and 1k resource units, respectively. In the *constant resource* condition, the resource value is fixed, whereas in the *variable resource* condition the population size is directly affected by the fixed resource increment 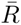. For sake of representation, 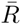 is not added to the resource values on the plot. B: Average *S_i_* and *R_i_* values at the last stage of the simulation (stage 60,000) in the constant (left) and variable (right) resource condition model. The horizontal dashed lines represent the expected *S_i_* and *R_i_* values for for the portion of the genome that is not subject to selection, used as a control region, indicated by *N_i_* in our model. Mutation rate is set to 0.001 per site and recombination rate is set to 0.01. *t_m_* corresponds to the age of sexual maturity.

Population genetic variance for *S_i_* reaches low values from stage 0 until about sexual maturation, and has its minimum around sexual maturation also for *R_i_,* as a consequence of strong purifying selection. After the onset of sexual maturation, genetic variance for both *S_i_* and *R_i_* increases rapidly, reflecting weakened purifying selection at later ages (FIG. 4).

We tested the effects of resource availability on population size oscillation and genome evolution. Even with constant resources, population size fluctuates due to the increased overall mortality induced by population size exceeding available resources (see II.b). In the *constant resource conditions,* population oscillations amplitudes were comparable when resources were set to 1k or 5k units (FIG. 5A). However, under *variable resource* conditions, population size is a function of the stage-dependent resources increment (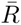) parameter. Population size oscillations had elevated amplitude in early stages, and stabilized to smaller amplitudes when 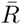 was high (5k units) (FIG. 5A). However, when 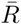 was set to 1k units (or lower), constraining population size to lowever values, population size oscillations did not stabilize, indicating that smaller populations are more unstable than larger populations. Importantly, in the *variable resource* condition, if 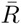 is large, resources do not oscillate with large amplitude once the population stabilizes (FIG. 5A). Decreased oscillation amplitudes in both population size and resource units is due to *S_i_* and *R_i_* values evolving towards a stable equilibrium (FIG. 5A). This was not observed when resources were variable and 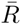 was small, in which case also the age-dependent distribution of *S_i_* in the post-reproduction phase was lower than in larger populations or in the *constant resource* condition (FIG. 5B), highlighting that in our model genome evolutionary dynamics not only depend on resource availability (constant or variable), but also on population size. Interestingly, we observed that smaller populations, in the *variable resource* condition, evolved to a status of increased early mortality, even before sexual maturation. This result was not a rare event and emerged in all our simulations. Additionally, in our model the age-dependent reproduction evolves in a different fashion from survival as – in the *variable resource* condition – *R_i_* have the same behavior in small and large populations (FIG. 5B, right). In fact, while *S_i_* drop right after *t_m_* in small populations under variable resources, *R_i_* dynamics are superimposable in large and small populations (FIG. 5B). This behavior might be explained by the fact that in our model only survival is penalised by a factor μ when population size becomes larger than available resources (see II.b).

### B. Asexual model

In the asexual model, we asked what the effects are of population size and availability of constant or variable resources on population dynamics and genome evolution (FIG. 6), and how these dynamics are different from the sexual model. In the *constant resource* condition, population oscillations decreased amplitude during long simulations (60k stages) both in larger and in smaller populations (FIG. 6A). Interestingly, unlike what we observed in the sexual model, in the *variable resource* condition, small-scale population size differences did not result in balanced equilibrium between resources and population size in the *variable resource* condition (FIG. 6A). However, we did observe an increase in the oscillation period between early and late stages of the simulation, as later stages of the simulation have longer oscillation periods than earlier stages. Additionally, the genome of asexually reproducing populations, both in the constant and, to a lesser degree, in the *variable resource* condition, did not reach elevated values of *S_i_* and *R_i_* both in large and small populations – at least in the tested population-size ranges – suggesting that, in populations reproducing asexually, selection for increased early survival and reproduction was weaker or had a slower evolution than in populations reproducing sexually (FIG. 6B).

**FIG. 6.**
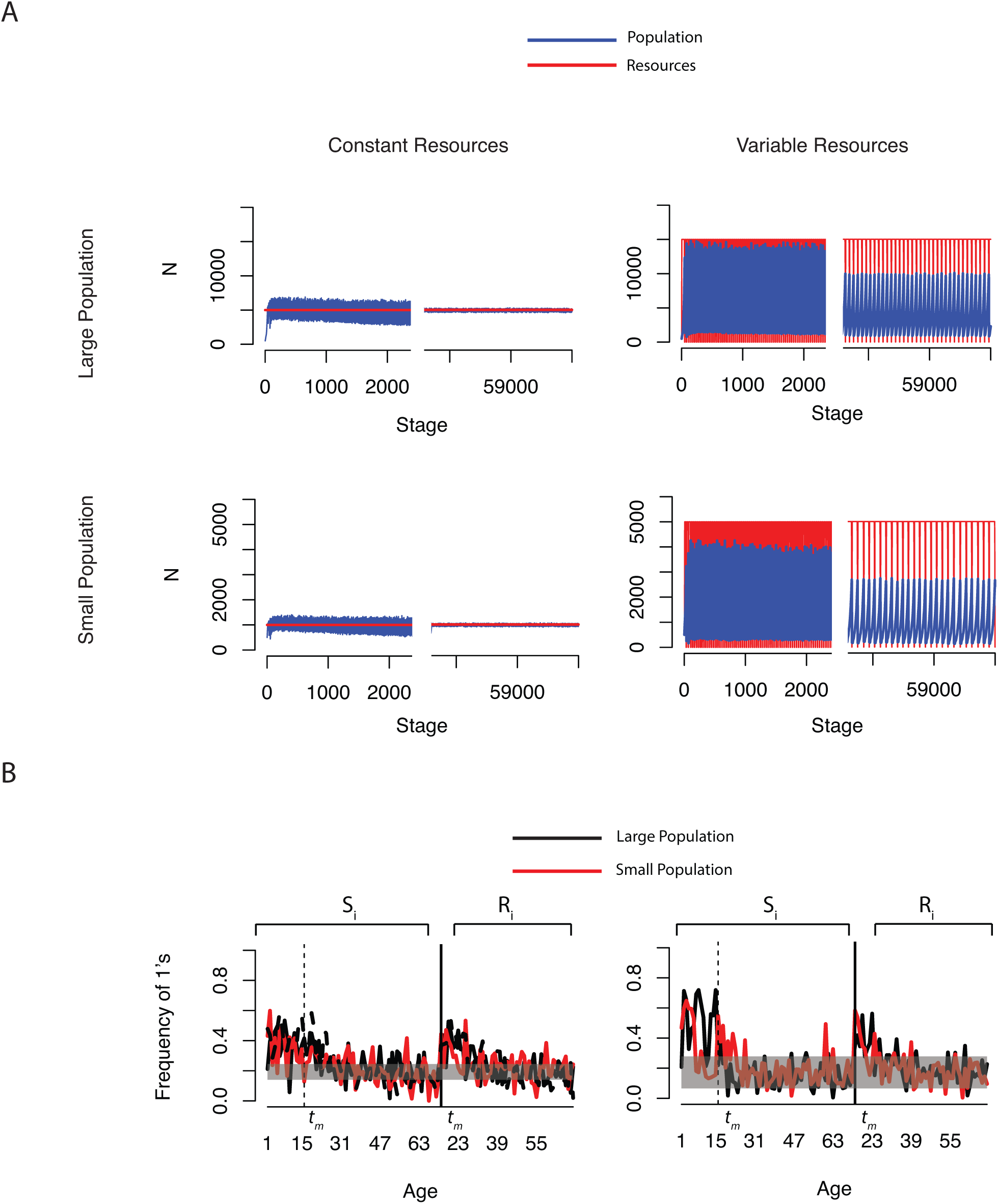
Population, resources and genomic dynamics at different population sizes in the asexual model. A: Population and resource dynamics in larger and smaller populations, under constant (left) or variable (right) resource conditions. Population and resource values are displayed on the y axis. Resources are shown in red and population in blue. Each plot is split in two parts, with early stages on the left, and late stages on the right. Each run consists of 60,000 stages. Large and small populations are determined by the resource value, set to 5k and 1k resource units, respectively. In the *constant resource* condition, the resource value is fixed, whereas in the *variable resource* condition the population size is defined by the fixed resource increment R. B: Average *S_i_* and *R_i_* values at the last stage of the simulation (stage 60,000) values in the constant (left) and variable (right) resource condition model for small and large populations. In the *constant resource* model we added a third plot (dashed black line) for *very large* populations, with 25k units of fixed resource. The horizontal shaded area represents the interval of values for *S_i_* and *R_i_* for the portion of the genome that is not subject to selection, used as a control region. Mutation rate is set to 0.001 per site and recombination rate is set to 0.01. *t_m_* corresponds to the age of sexual maturity.

### C. Survival and death rates

We calculated the observed death rate in both the sexual and asexual model, for the constant and variable resource model, in small and large populations (FIG. 7).

**FIG. 7.**
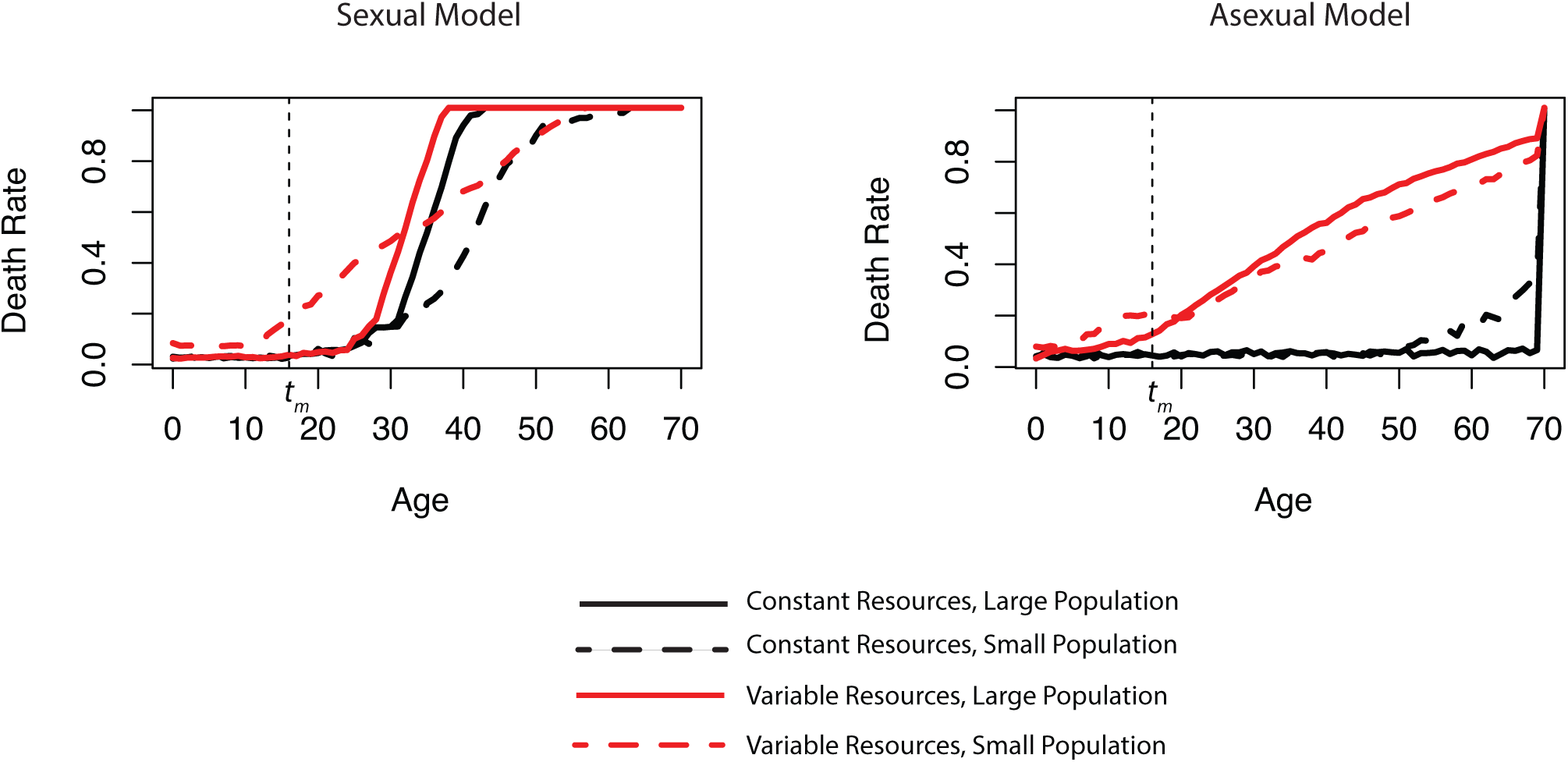
Observed death rate in the sexual and asexual models, with constant and variable resources, in large and small populations. Death rate is expressed, for each age *a* at stage *s*, as 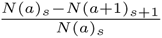, with *N(a)_s_* as the number of individuals of age *a* at stage *s*. The displayed values are the average death-rates for the last 100 stages of 60k stages-long simulations. *t_m_* corresponds to the age of sexual maturity.

Since in our model resources affect survival probabilities when the population size exceeds the available resources (see II.b), actual (i.e. observed) mortality at a specific time point is expected to differ from the mortality predicted solely by the distribution of the *s_k_* values.

In the sexual model, both in the *constant resource* condition and in the *variable resource* condition, once reached equilibrium between population and resources, the age-dependent death rate remained low before sexual maturation and increased after sexual maturation in large populations with slightly different dynamics, as in the *constant resource* condition the increase in mortality happened later than in the *variable resource* condition (FIG. 7). Small populations in the *variable resource* condition evolved high early mortality – i.e. before sexual maturation – but had an overall slower increase in the age-dependent death rate, as shown by the smaller slope in the death rate (FIG. 7).

In the asexual model, under constant resources, we find no age-dependent change in the rate of death both in large and small populations, except for an increase in death rate after age 50 in small populations (FIG. 7). This might result from the evolution of *S_i_* in the asexual model under constant resources, which does not significantly deviate from the expected values reached in the control genome region (FIG. 6B, left). However, the constant death rate across all the ages in the *constant resource* condition is associated with extemely high population oscillation stability, as indicated by the limited population size oscillation amplitude (FIG. 6A). Interestingly, we do observe an age-dependent increase in the rate of death in both the small and large populations for the *variable resource* model (FIG 7). This behaviour is associated with high amplitude in population size oscillation, which in our model leads to increased overall mortality (FIG. 6A).

## IV. Discussion

Studying the changes in individual probability of survival and reproduction is the focus of gerontology, experimental aging research and evolutionary biology. Aging research is ultimately focused on identifying the molecular mechanisms that affect the changes in the probabilities of surviving and reproducing throughout one individual’s lifetime. On the other hand, evolutionary biology – and in particular population genetics – is interested in the genetic changes that affect the probabilities of survival and reproduction over several generations, as they directly determine biological fitness, i.e. how likely it is to transmit specific alleles from one generation to the next.

In our model, we combine the “biology of aging” and the “evolutionary biology” perspectives on survival and reproduction and develop a novel *in silico* system where distinct genomic elements separately determine the probabilities of surviving or reproducing at a specific life stage. Using this approach, we can investigate how natural selection molds the allele frequencies that determine survival and reproduction at different life stages within the lifetime of an individual – as well as throughout generations – in the population’s gene pool.

We let populations evolve either sexually or asexually, under the effect of mutation and recombination (in the sexually reproducing populations). As populations evolve under fixed or variable resource conditions, we monitor population and genomic dynamics throughout the simulation. In particular, we monitor the evolution of differential survival and reproduction probabilities at the different individual ages allowed by our model. We compare the age-dependent values that survival and reproduction probabilities reach throughout the simulation with those achieved in a neutral portion of the genome that is not subject to selection. This enables us to quantify the strength of selection, and to identify the portions of the genome that are under stronger selection, i.e. those that deviate from the neutrally evolving parts of the genome. We find that populations reproducing sexually evolve gene-pools that have high survival probabilities before sexual maturation and progressively lower survival probabilities after sexual maturation. This is in line with the expectation that early-acting deleterious mutations are efficiently removed by selection from the gene pool^9^. Importantly, genomic variance for genetic elements affecting survival is lower in the pre sexual-maturation phase – indicated by *t_m_* throughout the paper – indicating strong purifying selection in early life stages. Additionally, the genetic elements affecting early survival accumulate high levels of beneficial mutations. Importantly, in our simulation the probabilities to survive evolve low values after sexual maturation and the genetic variance for survival (as well as for reproduction), increases dramatically after sexual maturation, strongly indicating that the age-dependent decreased probability of surviving and reproducing, i.e. aging, is directly correlated with increased genetic variance, which results from the relaxed purifying selection to maintain elevated survival and reproduction in late-life stages. This result suggests that genes or gene variants affecting late-life survival and reproduction are expected to be associated with higher population variance compared to genes affecting early-life phenotyes, such as development and early reproduction.

In the sexual model, populations evolving under variable resources undergo large oscillations in population size during the first few thousand simulation stages. However, at later stages, oscillations in population size and resources decrease in amplitude and reach an equilibrium (FIG. 5A). Additionally, we provided environmental perturbations to populations “in equilibrium” by stochastically imposing 90% mortality to the whole population. This had no effect on population size oscillation (data not shown), i.e. the population-resource equilibrium re-established rapidly after the perturbation. Once reached this stable equilibrium between resources and population size oscillations, the *variable resource* model behaved like the *constant-resource* model. In future studies we will investigate how varying the *fixed resource increment* 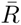 affects population oscillations and genome composition. Intriguingly, we observed that population size affects the probability to reach a resource-population equilibrium in the *variable resource* model, as smaller populations do not reach such balance. Larger populations display therefore higher stability and genetic plasticity compared to smaller populations. Interestingly, we find that sexually reproducing populations under *variable resource* conditions develop increased early-life *infant* mortality, i.e. the survival probabilities in the ages preceding sexual maturation significantly drop under two conditions: i) when population size is small and ii) in early stages of large population simulations, before the population-resource equilibrium is reached (FIG. 5B and FIG 7). Elevated initial *infant* mortality is associated with population instability and disappears when population and resource oscillations eventually balance each other. We find the increase in early mortality paradoxical, since the early ages preceeding sexual maturation, which in our model starts at age 16, should be under equally strong purifying selection, and any deleterious mutation affecting survival probabilities in this life stage should be rapidly removed from the gene pool. At the same time, we are aware that many species – including our own – display elevated infant death rates, which then decline to a minimum around puberty. After puberty, death rate starts raising, as already observed by R.A. Fisher^3^. A possible interpretation for this surprising elevated *infant mortality* is that, in the *variable resource* condition, large population oscillations, characterized by continuous population contractions and expansions, weakens the action of selection to efficiently remove deleterious mutations acting at early ages. Additionally, deleterious mutations become more likely to reach high frequency in the population due to continuous bottlenecks^7^. Alternatively, it could be speculated that a potentially active mechanism could favour the fixation of early-acting deleterious mutations to buffer large population oscillations by lowering the population growth rate, ultimately benefiting population stability. However, these hypotheses need to be rigorously tested in future studies.

Unlike populations reproducing sexually, asexually reproducing populations under *constant resource* conditions do not evolve in our model an age-dependent differentiation in the genomically-encoded probabilities to survive and reproduce. In particular, we did not observe higher probability of surviving or reproducing in early life stages, compared to late life stages. This phenomenon can be observed at the genomic level, where the genomically-encoded survival and reproduction probabilities do not dramatically differ before and after sexual maturation (FIG. 6B). Similarly to what seen in the *S_i_*, the measured death-rates show – particularly in the *constant resource* condition – no variation (i.e. do not increase) with age (FIG. 7). Since we define aging as an age-dependent increase in the probability of death and as a reduction in the probability of reproduction, asexually reproducing populations – particularly under *constant resource* conditions – seem to not undergo aging. In our model, asexually reproducing populations under fixed resource conditions are associated to the absence of senescence. This phenomenon, which spontaneously emerged in our simulation, is surprisingly in line with the evidences supporting extreme longevity and the lack of senescence in several organisms that reproduce clonally^4^. However, some asexually reproducing species are also reported to undergo senescence^8^. Interestingly, our model shows to some degree a detectable increase in the age-dependent death-rate in asexually reproducing populations under the *variable resources* condition (FIG. 7). In fact, both in large and small populations – using the same resource and population ranges used for the sexually reproducing populations – population size and resource units oscillations did not evolve to a status of equilibrium characterized by lower oscillation amplitude. This result can be interpreted as a lower stability of populations reproducing asexually, compared to populations reproducing sexually, under varying resource conditions. In future studies it will be important to explore how sexually and asexually reproducing populations compete against each other under different resource conditions.

### A. Conclusions

For aging to be considered an adaptation, we would expect a decreased genetic variance for the genetic modules that determine late-life decrease in survival and reproduction. In contrast with this expectation, we observed an age-dependent increased genetic variance following sexual maturation, which can be interpreted as decreased purifying selection in late-life phenotypes. The fitness effect of beneficial mutations acting in late-life is negligible compared to mutations affecting survival and reproduction at earlier ages. Even in our model, which does not constrain survival and reproduction to become higher or lower at different life-stages, individual fitness was not maximized, as survival and reproduction probabilities were elevated only in early life, before and around sexual maturation, and then decayed after sexual maturation, due to the effects of relaxed purifying selection and consequent mutation load on late-life acting genetic modules. This finding is in line with previous studies that predicted a declining force of natural selection throughout lifespan^1,5,9^. Since the genetic elements acting on survival and reproduction in late life-stages accumulated more deleterious mutations, compared to those responsible for survival and reproduction in earlier life-stages, we observed that a large portion of the genome that affected individual survival and reproduction at later ages was strongly associated with high genetic variance.

In our model, the genetic modules that determine the probabilities to survive and reproduce at different time stages are free to evolve due to mutation, recombination (in the sexual model) and selection. The simulations analyzed in this study were run at fixed recombination and mutation rates (indicated in the figure captions). Future studies will clarify how recombination and mutation rate optima evolve under different conditions and affect the age-dependent probabilities of survival and reproduction.

To note, our model does not allow for pleiotropic effects of genetic modules acting at different ages, since each genetic module is age-specific. We can hypothesize that in the presence of antagonistic pleiotropism, increased late-life mortality and decreased late-life reproductive potential could be potentially positively selected, granted that they provide a fitness advantage in early life stages.

Importantly, our model does not allow epistasis to take place in a direct genetic way, i.e. in our simulations there are no gene to gene interactions that lead to non-additive phenotypic effects, as each genetic module is independently responsible for its own phenotypic output. Although epistasis plays a key role in real biology, we deliberately did not implement it in our model for computational simplicity. The genetic modules for survival and reproduction that are implemented in our model, rather than being an *in silico* version of individual genes, can be understood as approximations of gene networks or pathways that influence either survival or reproduction at given ages. In this respect, epistasis might not be applicable to our model. Importantly, despite the absence of epistasis, our model reproduces several features of real population aging, such as increased early survival and rapid increase of the death rate and decrease of fertility after sexual maturation. Despite the absence of *sensu strictu* epistasis and pleiotropism, this model allows complex coevolution patterns between genetic modules that affect survival and reproduction at different life-stages, and provides an indirect test for the necessity of genetic epistasis and pleiotropism for the evolution of genetically-encoded biological aging. In other words, since our model evolves features typical of biological aging populations without implementing epistasis and pleiotropism, we can conclude that epistasis and pleiotropism – although important in organism aging processes – might not be necessary for the development of biological aging. Additionally, adopting a model that associates to individual ages discrete probabilities to survive and reproduce, enables to accurately calculate fitness at both individual and population level. Fitness measures factor in both survival and reproduction probabilities encoded in the genome, as well as observed survival and reproduction outputs. The study of how fitness varies throughout population evolution under different resource conditions and population size will be further developed in future studies. This model provides on one hand a powerful *in silico* system to generate hypotheses to test on real-life data, and on the other hand provides a numerical platform to test parameters derived from real biology, such as measured mutation and recombination rates, population size, resource availability and overall extrinsic risk of mortality. The use of this novel model to simulate the genome evolution in the context of biological aging opens new possibilities to understand how real populations evolve such a wide range of life history trait strategies.

## V. Authors Contribution and Acknowledgements

D.R.V. developed the model, contributed to the code and wrote the manuscript. A.S. wrote the python code and critically contributed to the improvement of the model.

We thank Fabio Iocco and Dmitri Petrov for the insightful suggestions and all the members of the Valenzano laboratory at the Max Planck Institute for Biology of Ageing for their constructive comments on the manuscript.

This work has been entirely supported by the Max Planck Society and by the Max Planck Institute for Biology of Ageing in Cologne, Germany.

